# Boosted cell-free gene expression for robust signal readout from a single-copy DNA template in microdroplets

**DOI:** 10.64898/2026.02.22.707295

**Authors:** Taro Furubayashi, Naohiro Terasaka, Kenya Tajima, Hiroyuki Noji

## Abstract

Cell-free gene expression in micro-compartments constitutes a chassis for biotechnology and synthetic biology. Protein synthesis from low concentrations of DNA, a single copy per compartment, is essential for in vitro evolution of biomolecules and synthetic cells. However, insufficient yield of protein synthesized from typically sub-picomolar DNA results in undetectable signals or inadequate activity of desired protein functions. Here we identify and largely mitigate yield-limiting bottlenecks of reconstituted in vitro transcription and translation (IVTT) at low DNA input. Systematic comparison of commercial reconstituted IVTT kits revealed that gene expression starts becoming limited by mRNA scarcity around 20-200 pM DNA input. We further uncovered that the standard ribosome concentration is excessive at low-DNA input and shortens the lifetime of translation. These findings led to a simple optimization recipe that combines supplementation with a highly active T7 RNA polymerase and a reduction in ribosome concentration, which synergistically amplified gene expression by ∼10-fold across diverse fluorescent proteins and enzymes. This low-DNA-optimized formulation in picoliter droplets achieved ∼94 nM protein expression from a single copy of DNA (∼0.12 pM). The user-friendly boosted IVTT protocol paves the way for straightforward functional screening and in vitro reconstitution of cellular functions in DNA-scarce environments.

**Graphical Abstract:** 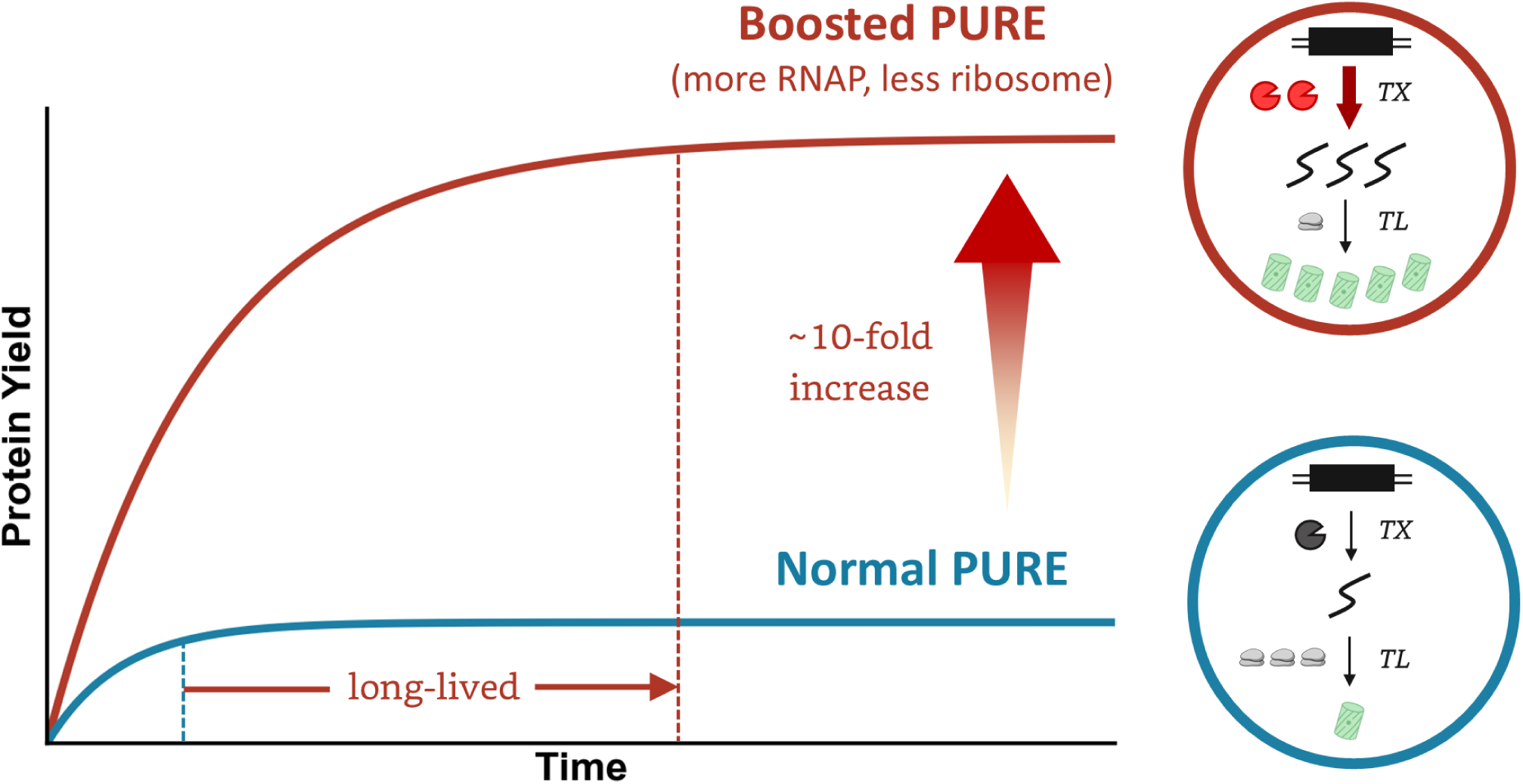

## Introduction

In vitro transcription and translation (IVTT) is a central biochemical platform widely used in synthetic biology and biotechnology^1–5^. IVTT decouples gene expression from the constraints of living cells while reproducing the essential information flow from genetic templates (DNA or RNA) to proteins. Applications of IVTT range from rapid prototyping of genetic circuits^6,7^, and metabolic pathways^8–10^ to molecular evolution^11–13^ and the bottom-up assembly of minimal synthetic cells^14–16^ to advance our understanding of complex living processes. Single-copy isolation of genetic variants is often required for in vitro evolution. Genotype–phenotype coupling by compartmentalization^17,18^ is the key concept, as it prevents functional crosstalk that would otherwise obscure which variants to select from a library containing active candidates and inactive “free-riders.” A typical strategy is to encapsulate each genetic variant in man-made micro-compartments^19,20^ such as water-in-oil droplets or lipid vesicles. Because hit discovery depends strongly on library size, fluorescence-activated droplet sorting (FADS) with droplet microfluidics^21,22^ is a major method to achieve high-throughput screening of typically 10^6^ to 10^7^ droplets in a day.

Low yields of gene products synthesized from a single DNA copy are a major hurdle in this context, as fluorescent signals often fall below the background noise^22,23^. Droplet size for FADS is typically ∼30 µm or larger in diameter, wherein a single DNA copy corresponds to <0.12 pM. Microarray-based screening^24^ or size-based fractionation of polydisperse droplets to enrich smaller compartments^25^ can circumvent this limitation by using sub-picoliter volumes to raise effective concentrations, but these approaches reduce throughput and add technical complexity. Another approach is DNA amplification before or during IVTT^26,27^, but amplification is inherently noisy and can obscure the true activities of genes under screening. Moreover, integrating DNA amplification with IVTT is challenging because the two reactions require different optimal buffer conditions^26,28^, often necessitating separate amplification and IVTT steps.

The PURE system^29,30^, a reconstituted system with highly purified components, is now a standard IVTT platform for this dilute single-copy DNA regime. Its high purity enables IVTT from a single DNA copy, which can be degraded by nucleases in crude cell extract-based systems^6,31,32^. Owing to the well-defined composition, PURE IVTT also provides a tractable framework for systematic analysis and engineering of reconstituted gene expression in artificial environments. Previous optimization efforts of the PURE system^33,34^ for higher protein yields focused only on high-DNA conditions, which may not be suitable for low-DNA conditions. Experimental and theoretical studies^35–37^ have shown that predicting PURE dynamics remains challenging due to nonlinear interactions among dozens to hundreds of molecular components. Therefore, a deeper understanding of PURE is essential for further development and more advanced applications.

We reasoned that at low DNA input, reaction bottlenecks in PURE IVTT may differ fundamentally from high-DNA conditions. In particular, we hypothesized that mRNA becomes scarce relative to translation components such as ribosomes, thereby altering the optimal conditions. Guided by this idea, we set out to interrogate how DNA input reshapes the limiting factors of PURE IVTT. With updated understanding of DNA-scarce conditions, we further sought a simple optimization recipe for enhanced IVTT yield and demonstrate its applicability in picoliter droplets for DNA amplification-free workflows.

## Results

### DNA concentration-dependent gene expression profiles of commercial PURE systems

To characterize DNA-concentration-dependent gene expression dynamics of the PURE system, we first titrated linear DNA templates encoding mNeonGreen (mNG) under the T7 promoter using three commercial PURE formulations: PUREfrex1.0, PUREfrex2.0 and PURExpress. PUREfrex2.0 is an updated formulation of PUREfrex1.0 for higher protein yield at high DNA input. PURExpress was reported to produce comparable yield of YFP as PUREfrex2.0 at high DNA input, but gene expression lifespan was shorter with mRNA degradation^38^. The DNA concentrations ranged from 0.2 to 2000 pM, and endpoint fluorescence at 15 h was used as a proxy for protein yield (Figure 1a). At 2000 and 200 pM DNA input, PUREfrex2.0 exhibited the highest protein yields, followed by PURExpress. A DNA input of 200 pM already saturated protein yield in PUREfrex1.0 and PURExpress. PUREfrex1.0 was unique in that DNA inputs from 2000 to 20 pM produced comparable protein yields. At 20 pM and below, all three PURE systems showed a DNA input-dependent decrease in protein yield, and PUREfrex1.0 performed best at sub-pM DNA conditions. PURExpress became least productive with low-DNA inputs (2 and 0.2 pM). Based on these profiles, we define the high-DNA regime as ≥200 pM DNA inputs and the low-DNA regime as ≤20 pM DNA inputs. End-point protein yield is largely insensitive to DNA input in the high-DNA regime, but becomes increasingly dependent on DNA input in the “transition band” between 20 and 200 pM.

**Figure 1.**
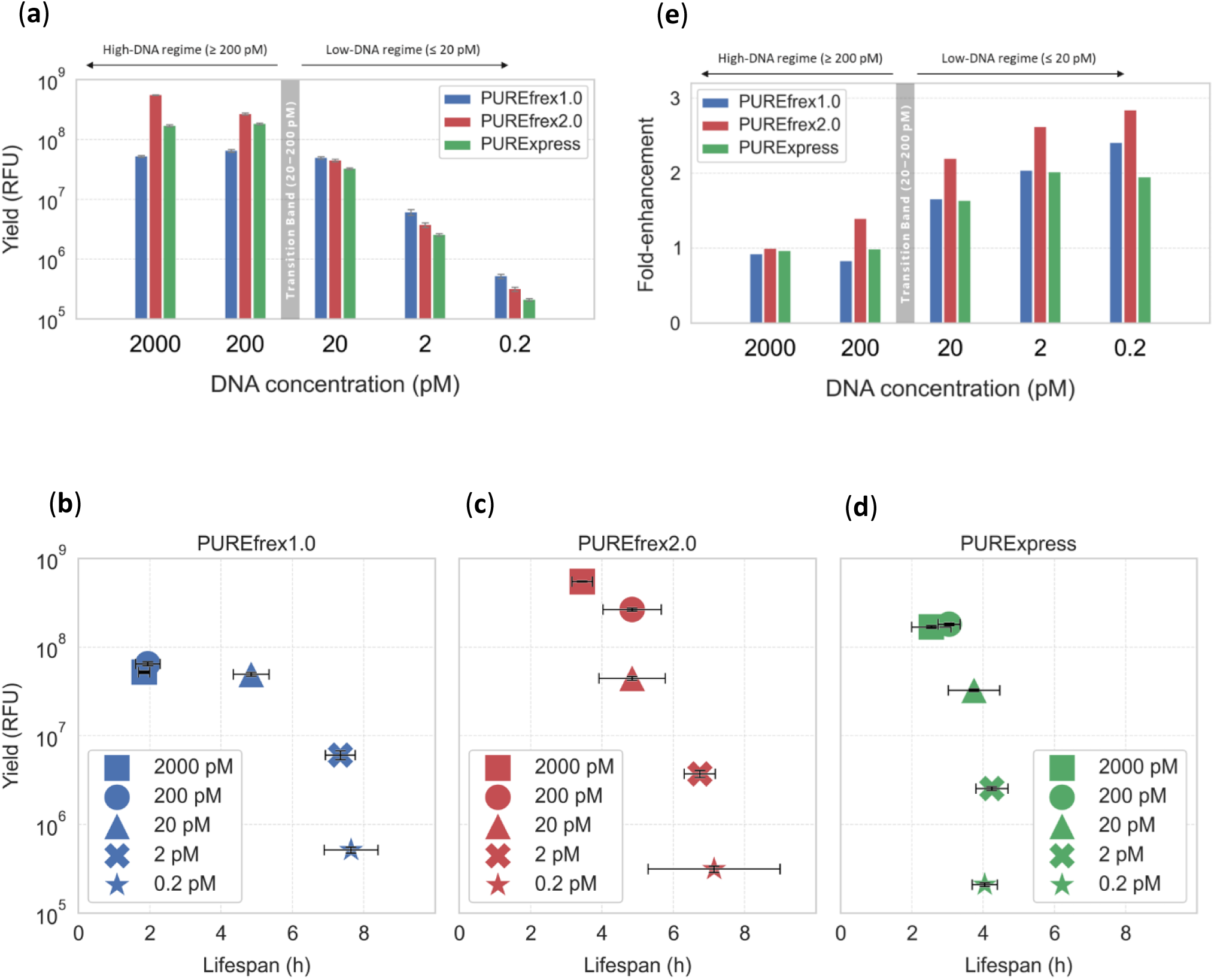
DNA concentration-dependent mNG expression profiles of commercial PURE systems. **(a)** mNeonGreen (mNG) yields as endpoint relative fluorescence units (RFU) at 15 h of PUREfrex1.0, PUREfrex2.0, and PURExpress with varied DNA concentration. Error bars indicate standard deviations (n = 3). (**b-d**) Yield-lifespan plots for (**b**) PUREfrex1.0, (**c**) PUREfrex2.0, and (**d**) PURExpress. Symbols denote input DNA concentration. Error bars indicate standard deviations (n = 3). Lifespan is defined as the time point at which mNG expression rate decreased to 30% of its maximum. (**e**) Fold-enhancement of IVTT levels as the ratios of mNG yields with and without supplementation of wild-type T7 RNA polymerase at 10% (v/v). The transition band regimes (20-200 pM DNA) are shown as gray bars in the panel (a) and (e).

Yield–lifespan analyses (Figure 1b–d) and kinetic profiles of gene expression (Figure S1) revealed distinct behaviors among the three PURE systems depending on DNA input concentrations. Here we defined lifespan as the time point at which mNG expression rate decreased to 30% of its maximum. In the high-DNA regime, PUREfrex1.0 was the shortest-lived, whereas PUREfrex2.0 was substantially longer-lived. Remarkably, in the low-DNA regime, lifespans of PUREfrex1.0 and PUREfrex2.0 were significantly prolonged, while PURExpress stayed short-lived compared to PUREfrex1.0 and PUREfrex2.0.

We postulated that augmenting TX by supplying additional T7 RNA polymerase may increase protein yield in the low-DNA regime, where mRNA scarcity rather than translation resources is expected to limit output. Indeed, the preferential increase in yield under low-DNA conditions supported this hypothesis (Figure 1e). Because the fold-enhancement was the lowest in PURExpress, we excluded PURExpress as a candidate for further optimization in the low-DNA regime. Together, these results indicate that PURE becomes transcription-limited under low-DNA conditions.

### Perturbation of transcription and translation with additional T7 RNA polymerase and ribosomes

Improvement of gene expression by supplementation of T7 RNA polymerase suggested that augmentation of mRNA production might be a promising solution for improved protein production in low-DNA conditions. Notably, the benefit of increased mRNA production emerged across the 20–200 pM transition band. Therefore, we designed systematic perturbation experiments of transcription (TX) and translation (TL) by titrating additional T7 RNA polymerase and ribosomes into PUREfrex1.0 and PUREfrex2.0 at 20 and 0.2 pM DNA.

To find a potent transcription enhancer, we first sought highly active T7 RNA polymerase variants that could enhance IVTT under DNA-limited conditions. We quantified mNG output after supplementing IVTT mixtures with wild-type or engineered T7 RNA polymerase variants from several suppliers. We found that T7 RNA Polymerase ver.2 (hereafter T7Pol-v2) from TaKaRa and the T7 RiboMAX™ Express system from Promega improved protein yields better than the wild-type T7 RNA polymerase (Figures S2 and S3). Considering cost efficiency, we used T7Pol-v2 to augment TX in all subsequent analyses.

At 20 pM DNA, TX augmentation with T7Pol-v2 consistently increased mNG production in both PUREfrex formulations (Figure 2a-b). Endpoint fluorescence rose with increasing T7Pol-v2 up to 15% (v/v). In contrast, supplementing ribosomes above the standard concentrations (i.e. 1000 nM for PUREfrex1.0 and 2000 nM for PUREfrex2.0) tended to reduce yields. Notably, however, co-supplementation with T7Pol-v2 and moderate ribosome supplementation in TX-augmented PUREfrex1.0 increased protein yields (Figure 2a), implying a subtle balance between mRNA-dependent increase and ribosome-dependent decrease of protein synthesis. PUREfrex2.0 was consistently inhibited by additional ribosomes, which may reflect more abundant translation machinery and reaction substrates than PUREfrex1.0. Under the titrated conditions at 20 pM DNA, the maximum protein yield was obtained with PUREfrex2.0.

**Figure 2.**
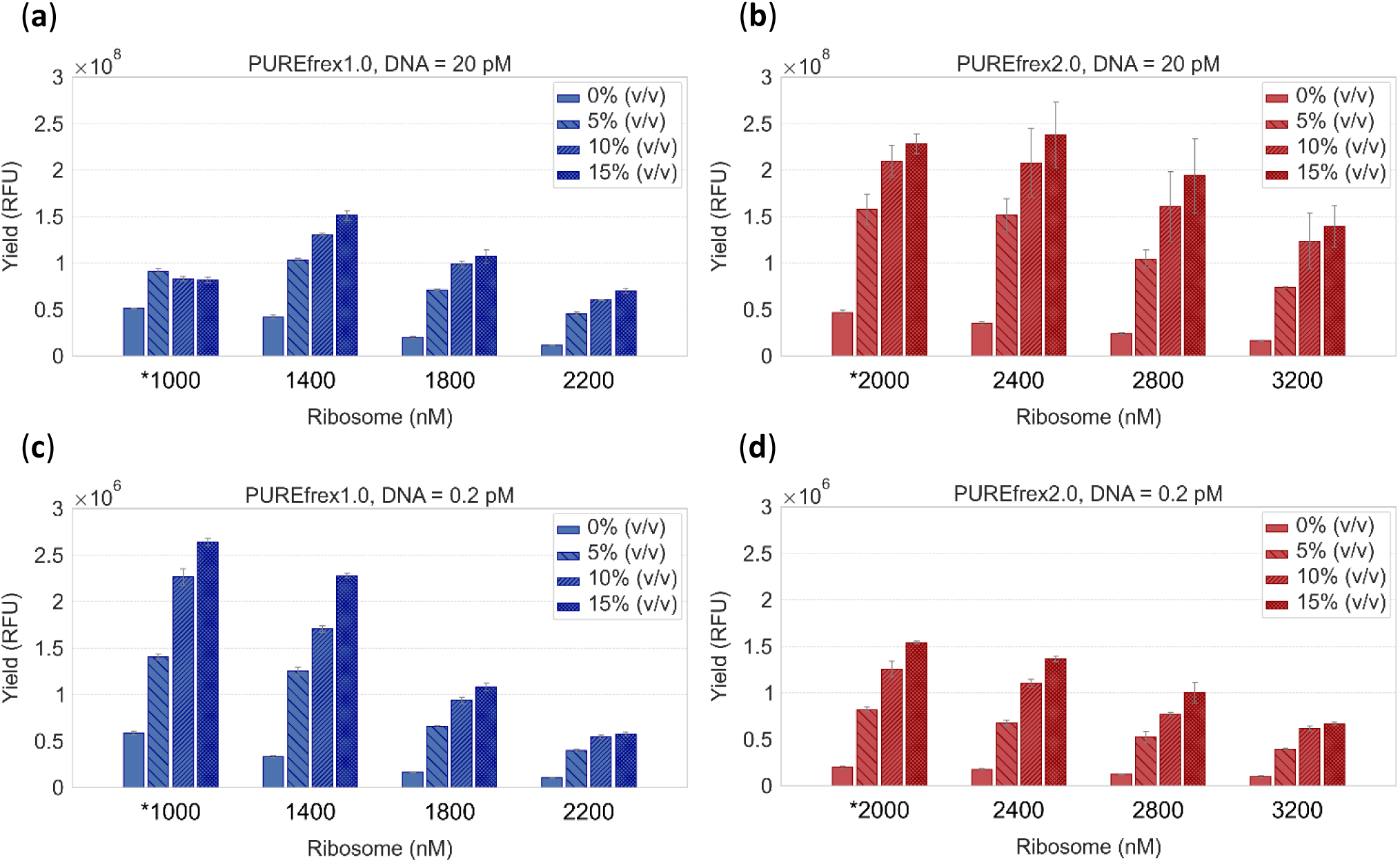
Effects of TXTL perturbations with additional T7 RNA polymerase ver.2 and ribosomes. mNG yields at 15 h for PUREfrex1.0 or PUREfrex2.0 at DNA inputs of 20 or 0.2 pM. (**a**) PUREfrex1.0 at 20 pM DNA (**b**) PUREfrex2.0 at 20 pM DNA (**c**) PUREfrex1.0 at 0.2 pM DNA (**d**) PUREfrex2.0 at 0.2 pM DNA. T7 RNA polymerase ver.2 was added at concentrations indicated in the legends. Starred values of ribosomes at the leftmost denote the standard concentrations specified in the manufacturer’s instructions. Error bars indicate standard deviations (n = 3).

At 0.2 pM of DNA, there was a consistent trend in TX and TL perturbation experiments. Additional T7Pol-v2 always led to higher protein synthesis, while more ribosomes were always detrimental in both PUREfrex1.0 and PUREfrex2.0 (Figure 2c-d). This is consistent with our “mRNA-limiting” hypothesis because mRNA concentration is still arguably lower compared to the translation machinery (e.g. ribosome concentrations are typically in µM orders in PURE systems). In this sub-pM DNA condition, PUREfrex1.0 exhibited higher yields than PUREfrex2.0.

### Reduced ribosome concentration prolongs active translation with low-DNA input

Upon the unexpected observation that additional ribosomes decreased mNG signals, we speculated that the optimal ribosome concentration might be lower than the manufacturer’s recommended level in the low-DNA regime. To identify the optimal ribosome concentration under TX-augmented conditions at 0.2 pM DNA (T7Pol-v2 = 15% (v/v)), we titrated ribosome concentration from the standard level down to 200 nM. To our surprise, ribosome reduction enhanced protein yields (Figure 3a) and markedly prolonged gene expression (Figure 3b-c). This effect was more pronounced in PUREfrex1.0 than in PUREfrex2.0. At 15 h, signal enhancement in PUREfrex1.0 saturated at 400 nM ribosomes, whereas PUREfrex2.0 produced similar signal levels between 1000 and 400 nM ribosomes. Even at 15 h, PUREfrex1.0 with 200 or 400 nM ribosomes remained translationally active (Figure 3b). These results support that lifespan- and yield-limiting factors differ between the high-DNA and low-DNA regimes, indicating a distinct optimization strategy is needed for improvement of protein expression from low-DNA. We further titrated potential IVTT enhancers (nuclease inhibitors, spermidine, EF-P, chaperones, and reducing agents) in the low-DNA-optimized condition (T7Pol-v2 = 15% (v/v), ribosomes = 400 nM), but none increased protein yields under the tested conditions (Figure S4).

**Figure 3.**
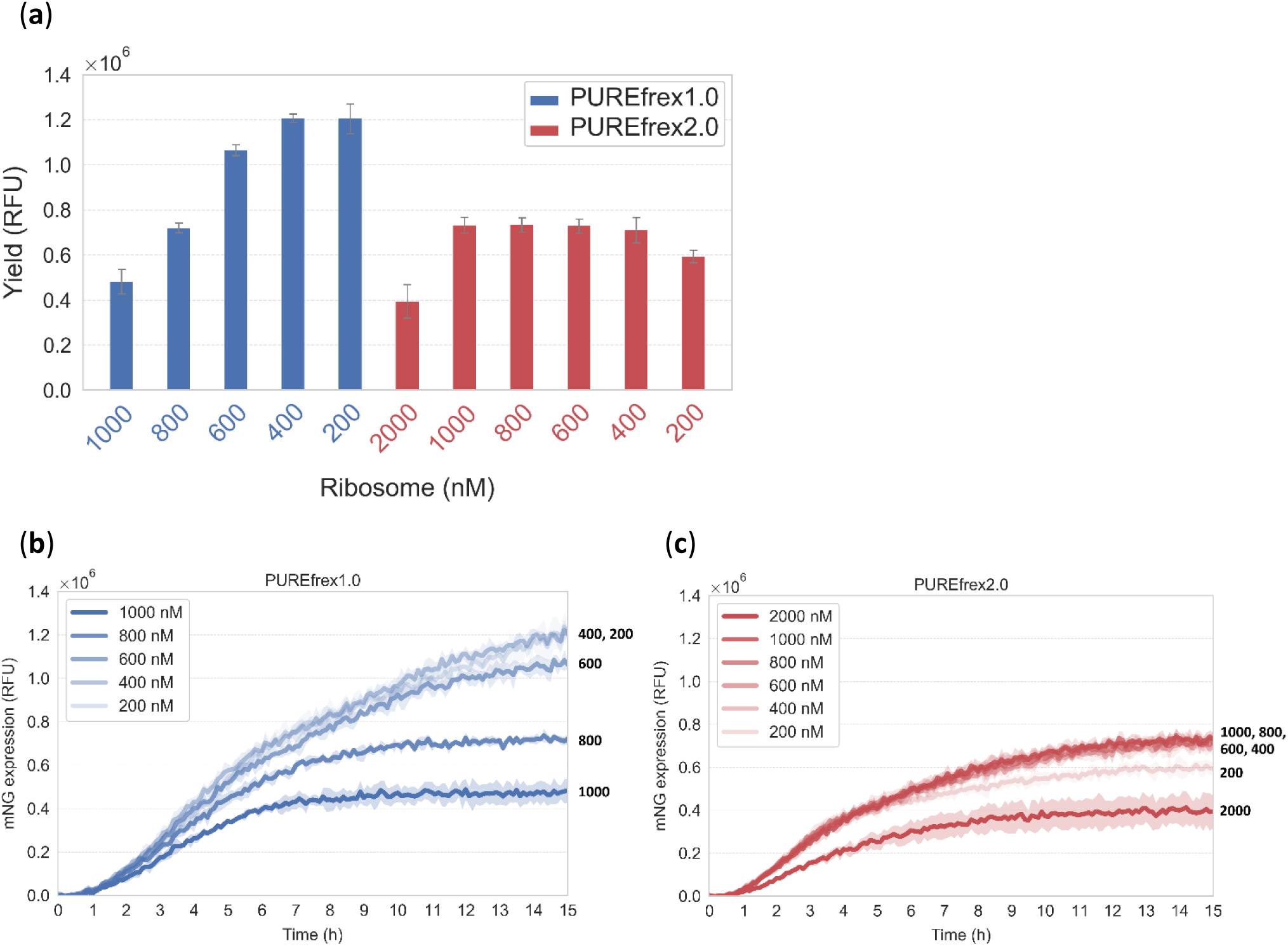
Effects of reduced ribosomes on mNG expression profiles in the low-DNA regime. mNG IVTT reactions were performed at DNA = 0.2 pM and T7Pol-v2 = 15% (v/v). (**a**) mNG yields at 15 h with reduced ribosome concentrations. (**b-c**) Kinetic profiles of mNG expression with reduced ribosome concentrations for (**b**) PUREfrex1.0 and (**c**) PUREfrex2.0. Numbers on the right ends of plots indicate the ribosome concentrations. Shaded areas indicate standard deviations (n = 3). Ribosome concentrations are shown in the legends.

### Enhanced gene expression and mechanistic insights under the low-DNA-optimized condition

We defined an optimal IVTT condition for 0.2 pM mNG DNA input: PUREfrex1.0 with T7Pol-v2 = 15% (v/v) and ribosomes = 400 nM (0.4× of the standard concentration). To assess the general applicability of this recipe, we assayed expression of several other proteins (mScarlet-I3, Savinase, NanoLuc, and ALP (*E. coli* alkaline phosphatase)) in bulk reactions (Figure 4a). We observed ∼10-fold signal amplification for most of the tested proteins compared to the standard PUREfrex1.0 condition. The similar extent of signal amplification across targets suggested a common underlying mechanism, most likely reflecting increased production of mRNA per DNA template. The only exception was ALP, which required buffer-exchanged T7Pol-v2 to a simple buffer to observe significantly enhanced IVTT signals of 6.2-fold (discussed in the Discussion section and Figure S5). To link TX and TL levels more directly, we quantified mNG mRNA and protein concentrations at 0, 3, 6, and 18 h (Figure 4b-d). We aimed to disentangle the contributions of TX augmented by T7Pol-v2 and ribosome reduction to changes in mRNA and protein levels. Endpoint protein yields showed that fold-enhancements in IVTT in ribosome-reduction-only, TX-augmentation-only, and TXTL optimized conditions were approximately 1.7-fold, 5.0-fold, and 10.8-fold, respectively (Figure 4b). The time courses showed distinct roles of T7Pol-v2 and ribosome reduction in increasing mRNA levels and prolonging the duration of translation, respectively (Figure 4c-d). Unexpectedly, ribosome reduction also increased mRNA levels at later time points (≥6 h) (Figure 4c). In summary, enhanced protein synthesis reflects synergy between elevated mRNA levels and increased protein output per mRNA (Figure S6), likely enabled by longer-lived translation (Figure 4e).

**Figure 4.**
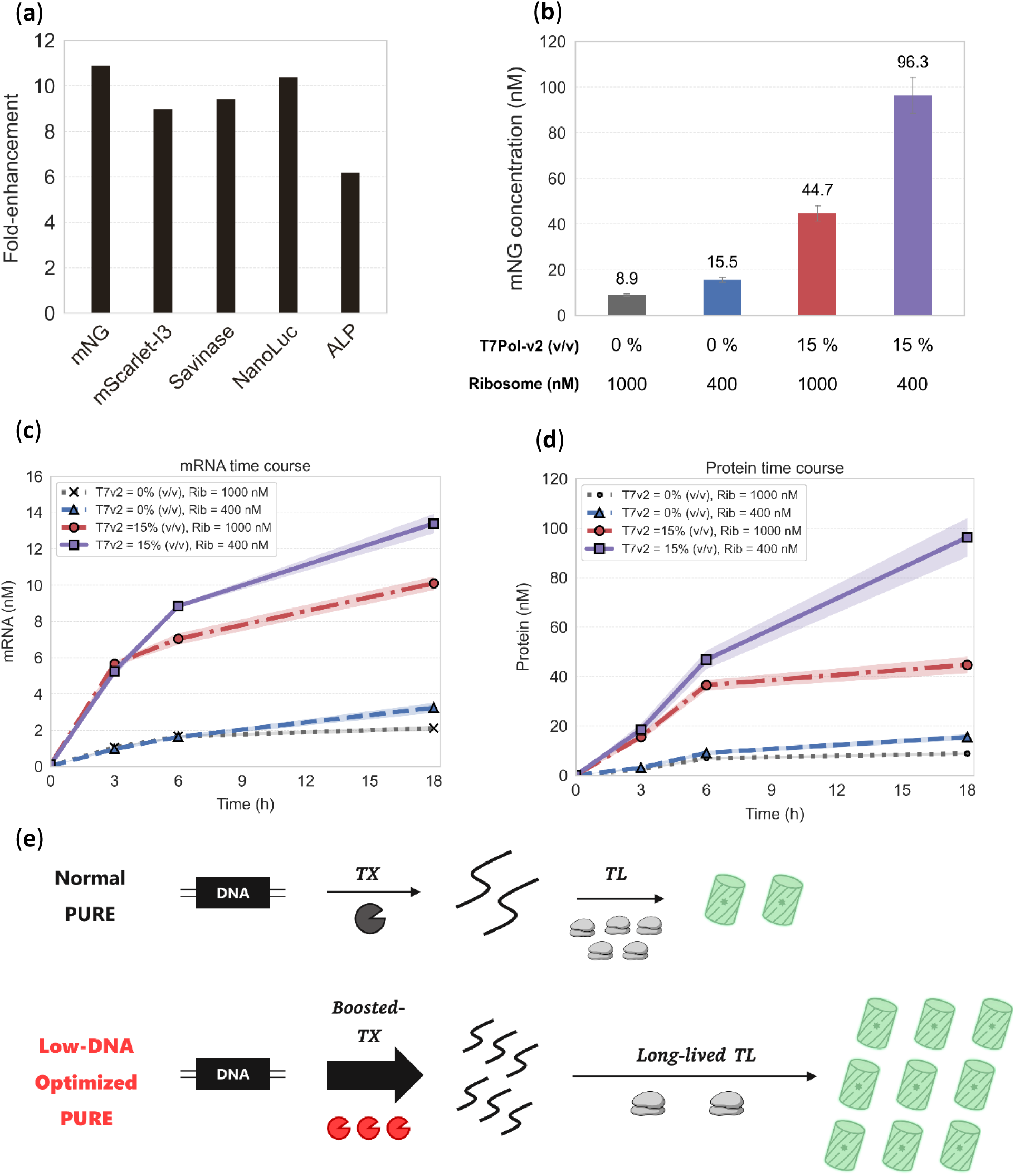
Enhancement of gene expression with the low-DNA-optimization and its expected mechanism. **(a)** Signal enhancement of fluorescent proteins and enzymes under the low-DNA-optimized IVTT condition. The ratios of yields with and without IVTT optimization are plotted. DNA inputs were 0.2 pM for mNG and 2 pM for the others. (**b**) Concentrations of mNG proteins at 18 h in PUREfrex1.0 IVTT under the normal, ribosome-reduced, TX-augmented, and optimized conditions. (**c-d**) Time courses of mRNA and protein of mNG in panel (**c**) and (**d**), respectively. T7Pol-v2 and ribosomes are indicated as T7v2 and Rib in the legends. (**e**) Schematic illustration of the expected mechanism of enhanced IVTT.

### IVTT in microdroplets and flow cytometry analysis for digital counting of DNA templates

To evaluate how the low-DNA-optimized PURE formulation improves gene-expression signals at the single-copy level, we encapsulated the optimized IVTT mixtures in monodisperse water-in-oil droplets of ∼30 µm diameter, corresponding to an effective concentration of ∼0.12 pM per DNA copy. Normal and low-DNA-optimized PUREfrex1.0 reactions supplemented with 0.1 pM mNG-DNA were compartmentalized such that each droplet contained 0 to a few DNA copies according to Poisson statistics. In the normal PUREfrex1.0 conditions, droplets containing a single DNA copy generated detectable but weak fluorescence signals (Figure 5a and b). Microscopy images showed only faint mNG expression, and fluorescence-activated droplet (FAD) analysis revealed digital peaks corresponding to 1, 2, and 3 DNA copies, yielding 19.1 nM, 38.9 nM, and 61.1 nM mNG, respectively (Figure 5c).

**Figure 5.**
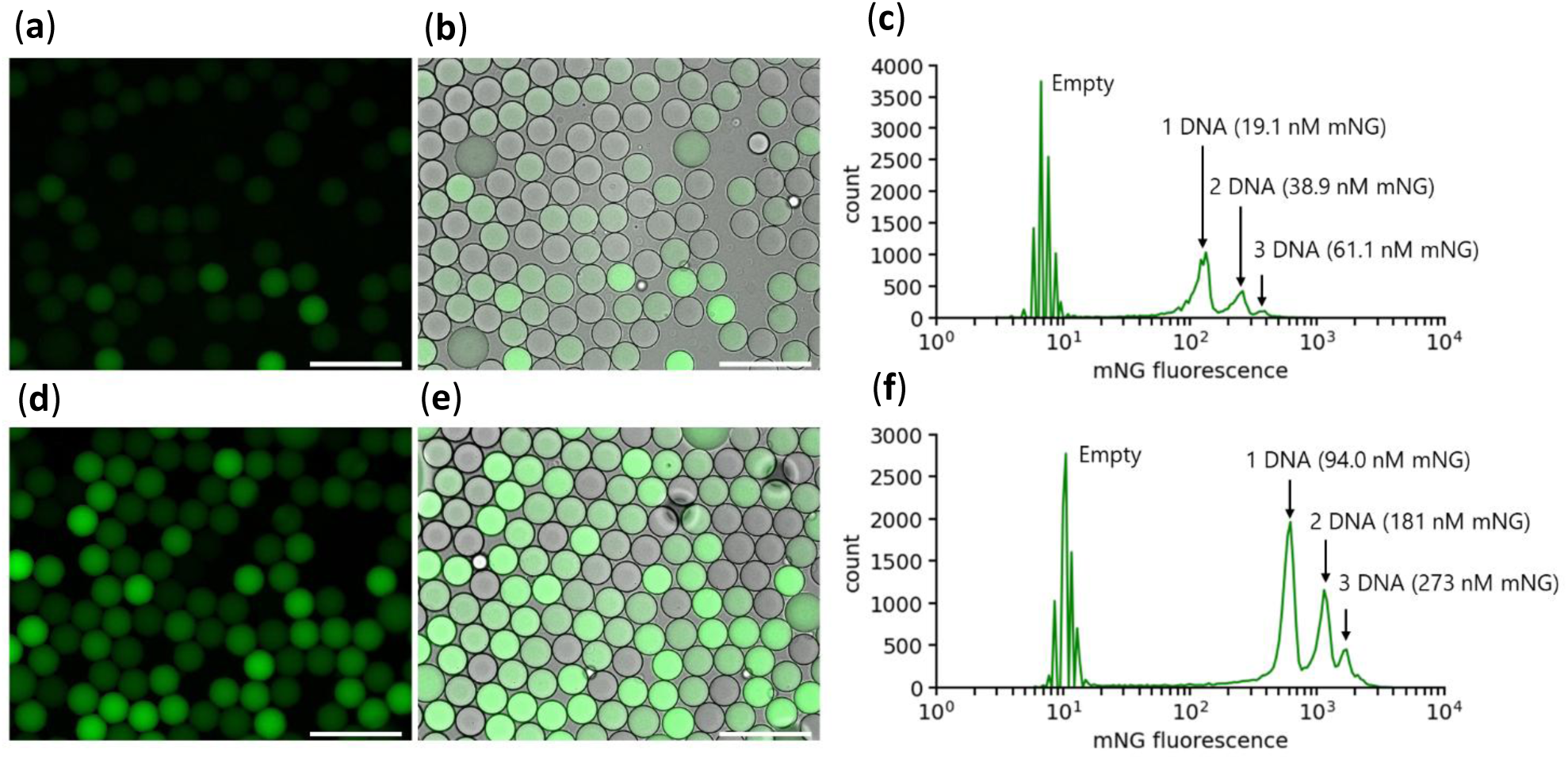
IVTT-containing microdroplets expressing mNG from single or multiple copies of DNA templates. (**a, d)** Fluorescent microscopy images of droplets expressing mNG. (**b, e**) Overlayed microscopy images of droplets expressing mNG (mNG fluorescence and DIC). (**c, f**) Fluorescence-activated droplet analysis of droplets expressing mNG. The amount of expressed mNG was quantified based on mean fluorescence intensity of each peak. Normal PUREfrex1.0 (**a-c**) or low-DNA-optimized PUREfrex1.0 (**d-f**) were used. Scale bars are 100 μm.

In contrast, the low-DNA-optimized PUREfrex1.0 produced stronger signals from a single copy of DNA. Microscopy images displayed clearly visible fluorescence (Figure 5d and e), and FAD analysis yielded distinct digital peaks corresponding to 94.0 nM, 181 nM, and 273 nM mNG for droplets containing 1, 2, and 3 DNA copies, respectively (Figure 5f). These values represent approximately a five-fold enhancement relative to the normal IVTT conditions, in line with the trend observed in bulk IVTT experiments. Thus, although digital counting is achievable with the normal PURE system, TX augmentation and ribosome reduction increase droplet fluorescence signals and facilitate robust discrimination of low-copy DNA templates in microdroplets without DNA amplification.

## Discussion

Protein yield from IVTT at low DNA input is a key determinant of the scope of advanced cell-free applications. In this work, we identified two bottlenecks that emerge in DNA-scarce regimes of PURE IVTT. First, transcription becomes yield-limiting due to mRNA scarcity relative to translation machinery as DNA input decreases through the 20-200 pM transition band. Second, an excess of ribosomes shortens the productive lifetime of translation, reducing final protein yield. Based on these insights, we developed a simple, user-friendly optimization recipe that increases protein yields by ∼10-fold in bulk reactions. Specifically, we measured ∼14 nM mRNA and ∼96 nM protein from 0.2 pM mNG-encoding DNA templates, corresponding to ∼70,000-fold (mRNA) and ∼480,000-fold (protein) increases in concentration relative to the input DNA concentration. These quantitative reference points provide a basis for condition-specific PURE formulation and tuning of in vitro artificial cells under limited DNA input.

Our results indicate a DNA-input-dependent shift in the yield-limiting step of PURE IVTT. Supplementation with T7 RNA polymerase had little effect on final protein yield under high-DNA conditions, whereas the response of PURE systems changed across the 20–200 pM transition band. PUREfrex2.0 was an exception: protein yield was not saturated at 200 pM and remained responsive to added T7 RNA polymerase at this condition, consistent with its higher translation capacity among the tested commercial PURE kits. Above this transition regime, mRNA availability is unlikely to be the limiting factor. Instead, translation factors, tRNAs, or energy supply are expected to determine final protein yield. This interpretation is consistent with many translation components in PURE being present in the order of 10 to 100 nM^33^, a range that can plausibly be saturated once mRNA supply is sufficiently high. In contrast, systematic perturbations of TX and TL at 20 and 0.2 pM DNA support the view that transcription becomes yield-limiting and that IVTT output in this low-DNA regime can be increased by TX augmentation with a high-activity T7 RNA polymerase variant. The similar fold-enhancement observed across several proteins suggests that the improvement is primarily driven by increased mRNA supply. Long lifespan of IVTT without mRNA degradation would be more influential in the low-DNA regime, and the limited response of PURExpress to TX augmentation may reflect reduced DNA template or mRNA availability at low DNA. The mRNA degradation in PURExpress probably due to residual nuclease activity was reported previously^38,39^, consistent with our results here.

Positive effects of reducing ribosome concentration at low DNA input were unexpected. Two effects were apparent: an extended period of productive translation and increased mRNA levels at later times. The former might be related to alleviated ribosomal traffic jams^40^ due to fewer ribosomes per mRNA, but this alone does not explain the latter. The late increase in mRNA may suggest that ribosomes waste GTP consumption because transcription also consumes GTP. It is known that excess ribosomes can engage translational GTPases in non-productive cycles, thereby accelerating futile GTP hydrolysis. For example, elongation factor-G (EF-G) can hydrolyze GTP in futile cycles on post-termination ribosomal complexes lacking ribosome recycling factor (RRF)^41^. In addition, initiation factor-2 (IF2) has been reported to undergo slow, uncoupled multiple-turnover GTP hydrolysis on ribosomes following 70S initiation complex formation^42^. If similar idling reactions occur in PURE, lowering ribosome abundance may suppress such futile GTP turnover and extend the functional lifetime of IVTT. The smaller benefit of ribosome reduction in PUREfrex2.0 is also compatible with this view, because PUREfrex2.0 is expected to harbor more translational GTPases than PUREfrex1.0 for higher translation capacity. Of note, however, it is unclear whether those futile reactions by GTPases really occur with apparently less substrates, such as mRNA and initiation or post-termination 70S ribosomal complexes. Quantifying GTP consumption dynamics is necessary to answer these questions.

We need to note that the presented low-DNA-optimized recipe may not directly apply to all types of proteins. *E. coli* alkaline phosphatase (ALP) provides a cautionary example, behaving differently from most tested proteins that showed approximately 10-fold signal enhancement. Under the same PURE conditions, ALP activity decreased as concentrations of T7Pol-v2 went up to 15% (v/v) (Figure S5a). We suspected that the stock buffer of T7Pol-v2 (composition not disclosed) is inhibitory to ALP, which may be more sensitive than the cytosolic monomeric proteins tested here. ALP is pH-sensitive and prefers alkaline conditions, and maturation requires disulfide bond formation and dimerization^43,44^ that normally occur in the periplasm. After exchanging the T7Pol-v2 storage buffer with a buffer containing 30 mM HEPES, pH 7.5, 100 mM potassium acetate and 1 mM TCEP, ALP activity increased to a 6.2-fold level, supporting our speculation (Figure S5b). This exception highlights the need to consider target-specific biochemical requirements and potential buffer incompatibilities when applying the recipe.

DNA amplification-free nature of the boosted PURE IVTT may benefit cell-free directed evolution campaigns with reduced screening noise stemming from stochastic nucleic-acid amplification^45^. Previous enzyme evolution with high-throughput fluorescence-activated droplet sorting (FADS) has usually relied on droplets in the tens of micrometers range, due to several technical limitations, which can make single-copy (sub-picomolar) templates difficult to detect. To circumvent this problem, a few studies have utilized intricate techniques such as in-droplet nucleic-acid amplification (e.g., in-droplet phi29-based DNA amplification^26^ or mRNA replication^46^). However, DNA amplification is highly stochastic, thus can blur true differences in enzymatic activity among gene variants due to variable copy numbers in compartments. The transcription-augmented PURE formulation in this study presents an alternative route for functional screening without introducing an additional amplification step that can inflate copy-number heterogeneity, and it may also reduce technical complexity and hands-on labor because it is implemented by simple reagent tuning.

Another potential advantage of PURE IVTT in droplet assays is reduced gene-expression variability relative to cellular expression, which may facilitate digital readouts. Noisy gene expression in cellular contexts is well known^47,48^ and arises from both intrinsic stochasticity and extrinsic sources such as growth state and plasmid copy number. In contrast, reconstituted PURE IVTT reduces cellular extrinsic fluctuations, so variability is expected to be dominated by intrinsic stochasticity in TX and TL. In our preliminary droplet experiments comparing digital mNG-DNA in optimized PURE with living *E. coli* single cells expressing mNG, the mean mNG signal levels were comparable, but the signal intensity distribution was markedly broader in *E. coli* containing droplets and digital peak separation was not achieved (Figure S7). Note that movement of *E. coli* cells in droplets is one of the potential reasons for the observation of broad peaks in FADS analysis. Further systematic analysis will be needed to precisely evaluate whether PURE offers a practical advantage for low-noise digital assays.

It is valuable to obtain high and reproducible protein output from sub-picomolar DNA templates with user-friendly modifications to commercial PURE systems. This optimization recipe can support straightforward droplet-based screening without additional nucleic-acid amplification, avoiding complicated signal-amplification workflows. This approach may also bring within reach previously inaccessible reconstitution of multi-subunit proteins and RNA–protein interactions that require near-micromolar components for their assemblies. In the context of bottom-up synthetic biology, it offers a way to maintain low DNA copy numbers for genotype–phenotype coupling while achieving protein levels sufficient for reconstituted functions such as genome replication, membrane remodeling, and metabolism. Together, these features make the low-DNA-optimized PURE a practical starting point toward novel cell-free systems that operate in DNA-scarce conditions.

## Acknowledgements

This work was supported by Japan Society for the Promotion of Science Grant-in-Aid for JSPS Fellows 21J01634 and 22KJ0540 (to T.F.), Grant-in-Aid for Early-Career Scientists 22K14794 (to T.F.), Transformative Research Areas (B) 21H05119 (to N.T. and T.F.), The Naito Foundation (to N.T.), Japan Society for the Promotion of Science Research Fellowships for Young Scientists 24KJ1042 (to K.T.), and Japan Science and Technology CREST JPMJCR19S4 (to H.N.).

## Author contributions

T.F. and H.N. designed the project. T.F. performed all the experiments and analyses except for droplet experiments. N.T. performed droplet experiments and analysis. K.T. expressed and purified proteins. T.F., N.T., K.T. and H.N. wrote the manuscript.

## Materials and Methods

### Linear dsDNA templates for IVTT reactions

DNA templates for IVTT reactions were synthesized by PCR using KOD One® PCR Master Mix (TOYOBO). Sequences of genes and primers used in PCR are listed in Supplementary Data. PCR products were purified by NucleoSpin® Gel and PCR Clean-up (MACHEREY-NAGEL) according to the manufacturer’s instructions. DNA was eluted with TE solution (pH = 8.0) (NIPPON GENE) and quantified by NanoDrop™ 2000c (Thermo Fisher Scientific).

### PURE IVTT reactions

IVTT reactions were performed using PUREfrex1.0 (GeneFrontier), PUREfrex2.0 (GeneFrontier), or PURExpress (New England Biolabs). Standard IVTT reaction mixtures were prepared according to the manufacturers’ manuals. All reactions were performed at 37 °C for 15 h unless otherwise specified. For TX perturbation in Figure 1e, T7 RNA Polymerase (New England Biolabs) was added to IVTT mixtures at 10% (v/v). For the perturbation experiments in Figure 2 and all subsequent optimized IVTT reactions, T7 RNA Polymerase ver.2 (TaKaRa) and *E. coli* ribosome (New England Biolabs) were used, and their concentrations are indicated in the corresponding figure legends.

Additional components other than T7 RNA Polymerase ver.2 (T7Pol-v2) were added to IVTT mixtures for enzyme activity assays in Figure 4a. For Savinase, BODIPY FL casein in EnzChek™ Protease Assay Kit (Thermo Fisher Scientific) was used as the reaction substrate. BODIPY FL casein was dissolved in Milli-Q water and added to the IVTT mixture at 0.02 mg/mL. For NanoLuc, IVTT reactions were performed without substrate, which was added immediately before signal measurement (details in the following section). For *E. coli* ALP, an ALP activation mixture was prepared with 80 µM DsbC (GeneFrontier), 20 mM GSSG (GeneFrontier), and 2 mM zinc acetate. The activation mixture was added to the ALP IVTT mixture at 5% (v/v). As the reaction substrate, FDP [Fluorescein diphosphate, tetraammonium salt] (Cosmo Bio) was added at 20 μM.

For signal enhancement of ALP IVTT in Figure S5b, buffer-exchanged T7Pol-v2 was used. 50 µL of T7Pol-v2 was mixed with 450 µL of an exchange buffer (the IDT duplex buffer supplemented with 1 mM TCEP) and loaded onto Amicon® Ultra Centrifugal Filter, 10 kDa MWCO (Millipore). The filter was centrifuged at 14,000×g for 10 min. 450 µL of the exchange buffer was loaded again on the filter and centrifuged at 14,000×g for 10 min again to recover 41 μL of buffer-exchanged T7Pol-v2. Concentration of T7Pol-v2 was measured by NanoDrop 2000c (Thermo Fisher Scientific).

### Signal measurements of IVTT reactions

Fluorescence and luminescence of PURE IVTT reactions were measured using a SpectraMax iD3 (Molecular Devices). Reaction mixtures were dispensed into 384-well Low Volume Black Round Bottom Polystyrene NBS Microplates (Corning) and sealed with Universal Optical Microplate Sealing Tape (Corning).

For fluorescence detection, excitation and emission wavelengths were 480 nm and 520 nm for mNG, Savinase, and ALP, and 548 nm and 592 nm for mScarlet-I3. Fluorescence was recorded every 6 min for 15 h at 37 °C, unless otherwise specified. Values shown in Figure 4a correspond to fluorescence at 15 h for mNG, mScarlet-I3, and Savinase, and at 5 h for ALP.

For NanoLuc luminescence, IVTT mixtures were incubated for 18 h at 37 °C and then diluted 100-fold with Nuclease-Free Duplex Buffer (IDT) to avoid detector saturation. Nano-Glo® Luciferase Assay Substrate (Promega) was used as the substrate. A substrate mixture was made by mixing 20 µL Nano-Glo substrate, 8 µL Recombinant Albumin Molecular Biology Grade (20 mg/mL; New England Biolabs), and 172 µL IDT duplex buffer. For measurement, 3 µL of the diluted IVTT mixture was combined with 27 µL of the substrate mixture. Luminescence at 460 nm was recorded at 37 °C for 6 min with a read interval of 90 s, and the average value was used for Figure 4a.

### In vitro transcription and purification of mRNA

mRNA encoding mNeonGreen (mNG) was synthesized by in vitro transcription (IVT) using a PCR product of a mNG gene (Supplementary Data). The IVT reaction mixture consisted of mNG-DNA at 2 nM, T7 RNA Polymerase ver.2 (TaKaRa) at 10 U/µL, T7 RNA Polymerase Buffer (TaKaRa), NTPs (Thermo Fisher Scientific) at 10 mM each, and Recombinant RNase Inhibitor (TaKaRa) at 1 U/µL. The IVT reaction was performed for 8 h at 37 °C. mRNA was purified using NucleoSpin® Gel and PCR Clean-up (MACHEREY-NAGEL) according to the manufacturer’s instructions. mRNA was eluted with Milli-Q water and quantified using NanoDrop™ 2000c (Thermo Fisher Scientific).

### Expression and purification of mNG proteins

**(a)** *E. coli* BL21-Gold(DE3) cells (Agilent Technologies) were transformed with pET-mNG. 2 L Erlenmeyer flasks containing 1 L LB medium with 100 μg/mL ampicillin were inoculated with 20 mL overnight cultures and incubated at 37 °C and at 150 rpm until OD600 reached 0.8. Protein expression was induced by adding IPTG (M&S TechnoSystems) to a final concentration of 0.4 mM. Cells were cultured at 25 °C overnight and then harvested by centrifugation at 5,000×g and 4 °C for 10 min. The cell pellet from one 1000-mL culture was resuspended in 40 mL LB medium, transferred to a 50 mL tube. The medium used for transfer was removed by centrifugation at 5,000×g and 4 °C for 10 min, decanted, and aliquots of the cell pellets were frozen in liquid nitrogen and stored at −80 °C until purification.

A cell pellet corresponding to 1000 mL of culture volume was resuspended in 10 mL of lysis buffer (50 mM sodium phosphate, 1 M NaCl, 20 mM imidazole at pH 7.4) and lysed by sonication (150 cycles of 1 sec on and 1 sec off with amplitude = 40, Q500 Sonicator, QSONICA). The soluble fraction in the lysate was recovered after centrifugation at 12,000×g and 4 °C for 10 min and loaded onto 1 mL slurry of Ni(II)-NTA resin (Nuvia IMAC resin, BIO-RAD) in a gravity flow column. After incubation for 30 min, the resin was washed with 10 mL of binding buffer (50 mM Tris-HCl, 300 mM NaCl, 20 mM imidazole at pH = 8.0) and eluted with 4 aliquots of 1 mL of elution buffer (50 mM Tris-HCl, 300 mM NaCl, 100, 200, 300, or 400 mM imidazole at pH 8.0). The eluted fractions were combined and dialyzed in 1 L of lysis buffer without imidazole at 4 °C twice. The protein quantity was determined by using Pierce 660-nm Protein Assay Reagent (Thermo Fisher Scientific) with NanoDrop OneC (Thermo Fisher Scientific). Final solution was frozen by liquid nitrogen and stored at −80 °C.

### Quantification of mNG mRNA and proteins in bulk IVTT

Concentrations of mRNA and protein during IVTT reactions in Figure 4c–d were quantified by reverse-transcription quantitative PCR (RT-qPCR) and fluorescence measurement using a SpectraMax iD3 (Molecular Devices). For mRNA quantification, One Step PrimeScript™ III RT-qPCR Mix (TaKaRa) was used. Absolute mRNA concentrations were estimated using a calibration curve generated from dilution series of in vitro transcribed mRNA. Time-sampled IVTT aliquots were diluted 500-fold with Nuclease-Free Duplex Buffer (IDT), incubated at 75 °C for 30 s, and placed on ice immediately before RT-qPCR. Elimination of DNA templates was not conducted because transcribed mRNA abundance was orders of magnitude higher than the DNA templates. For protein quantification, purified mNG protein was used to generate a calibration curve for fluorescence measurements. To minimize protein loss by adsorption, pipette tips were pre-coated with Recombinant Albumin Molecular Biology Grade (20 mg/mL; New England Biolabs) diluted 10-fold with IDT duplex buffer, and 1.5 mL PROKEEP protein low-binding tubes were used for protein dilution.

### Generation of water-in-oil microdroplet

The water-in-oil emulsion droplets were generated by On-chip Droplet Generator (On-Chip Biotechnologies) in 2D chip-800DG chips (1003002, On-Chip Biotechnologies) at 10 °C. The oil phase was prepared by mixing 008-FluoroSurfactant-5wtH (2003001, On-Chip Biotechnologies) and 008-FluoroSurfactant-0.1wtH (2003002, On-Chip Biotechnologies) at a 2:3 ratio. PURE system containing 100 fM of mNG-DNA, purified mNG (0 nM, 10 nM, 100 nM or 1 μM) or *E. coli* BL21-Gold(DE3) cells bearing pET-mNG cultured overnight at 25 °C after adding IPTG (OD600 = 0.015 in PBS) were used as the aqueous phase. Air pressures for the aqueous sample and the oil sample were set to 45 kPa and 50 kPa respectively.

### Microscopy imaging of droplet

The droplets containing cell-free translation mixture were collected in a microtube and incubated at 37 °C for 18 h to express proteins. The droplets were put into Cell Counting Slides (#1450015J, BioRad) and visualized on an ECHO Revolve microscope using FITC filter set (Ex, 470 nm; Em, 525 nm).

### Fluorescence-activated droplet analysis

The droplets containing cell-free translation mixture were collected in a microtube and incubated at 37 °C for 18 h to express proteins. The droplets were analyzed by On-chip Sort machine (On-Chip Biotechnologies) in 2D Chip-Z1001 chips (1002004, On-Chip Biotechnologies). The sheath fluid was 008-FluoroSurfactant-0.1wtH. The droplets bearing ∼30 μm diameter were gated based on the value of forward scatter and the histogram of mNG fluorescence (FL-2) was analyzed. In the histogram (FL-2), mean fluorescence intensity of droplets containing purified mNG (0 nM, 10 nM, 100 nM or 1 μM) was used to make standard curve for quantification.

---

**Figure S1.**
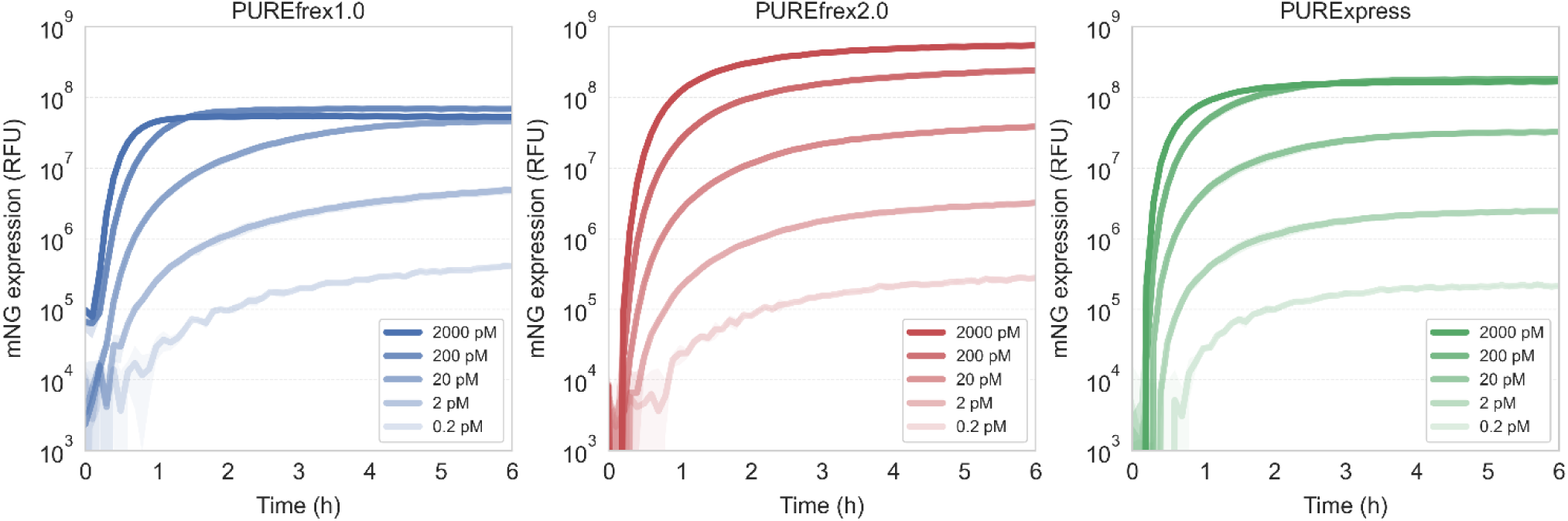
mNG expression kinetics of commercial PURE systems. Time courses of mNG expression (RFU) up to 6 h for PUREfrex1.0, PUREfrex2.0, and PURExpress with varied DNA concentration. Shaded areas indicate standard deviations (n = 3).

**Figure S2.**
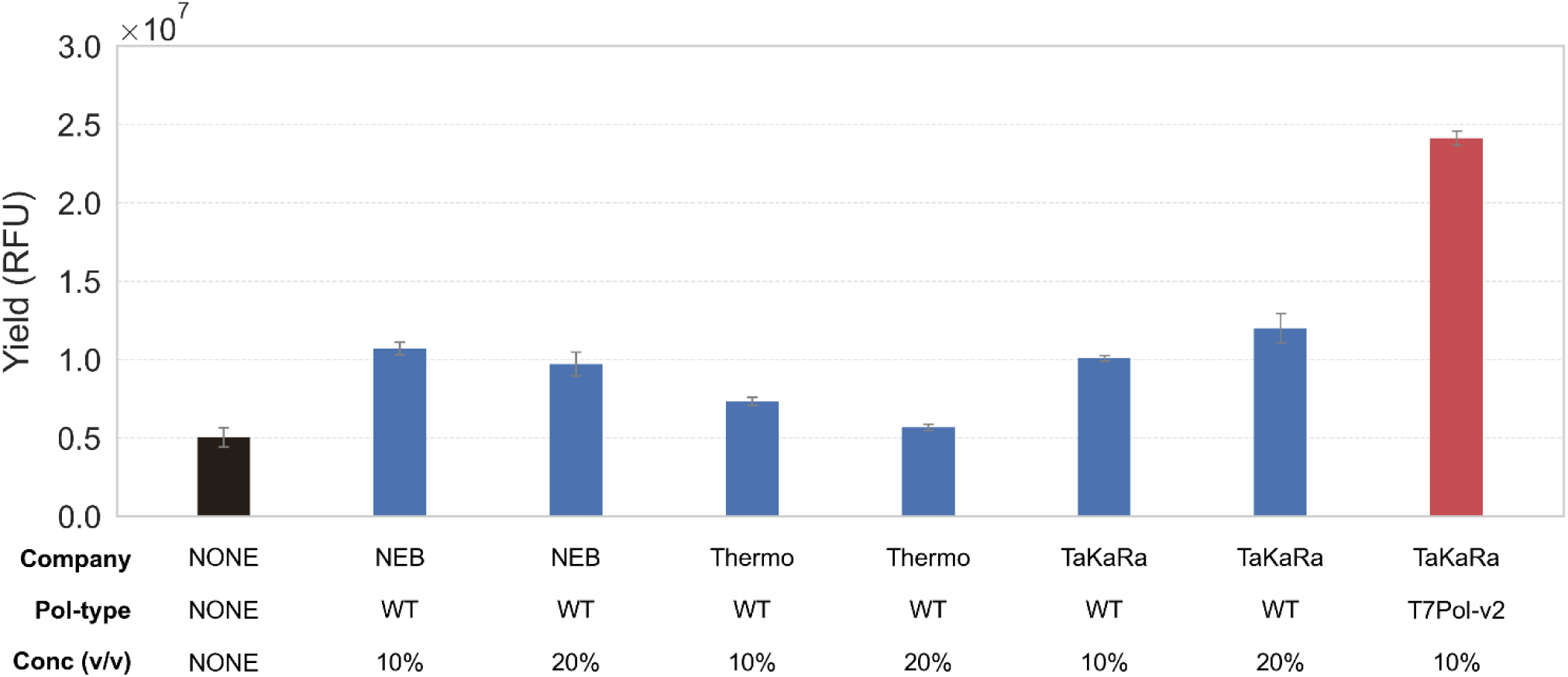
Comparison of commercial T7 RNA polymerases for enhanced IVTT. mNG yields at 15 h supplemented with commercial T7 RNA polymerases. Error bars indicate standard deviations (n = 3). PUREfrex2.0 was used with 2 pM DNA input. T7Pol-v2 is T7 RNA Polymerase ver.2 (TaKaRa). Black, blue, and red bars indicate the standard, wild-type T7Pol supplied, and T7Pol-v2-supplied IVTT conditions, respectively.

**Figure S3.**
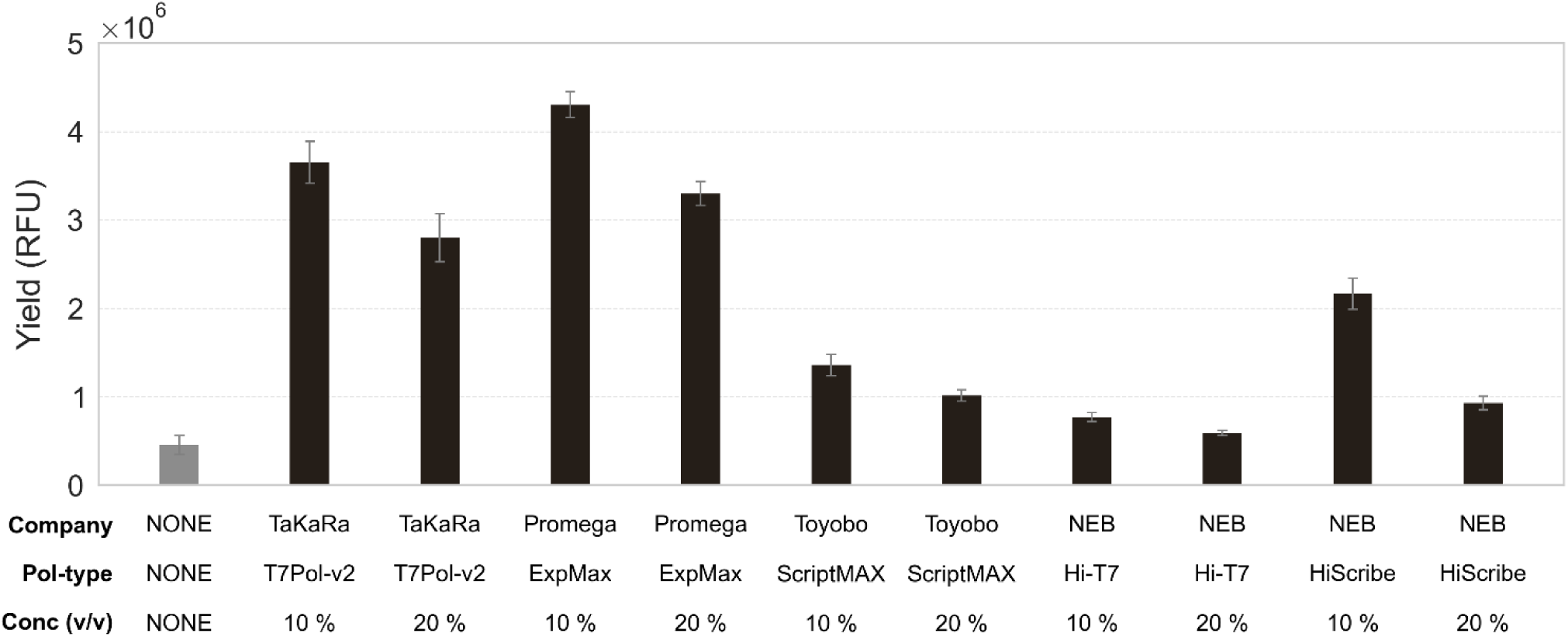
Comparison of commercial variants of T7 RNA polymerase for enhanced IVTT. mNG yields at 15 h supplemented with commercial variants of T7 RNA polymerases. Error bars indicate standard deviations (n = 3). PUREfrex2.0 was used with 0.2 pM DNA input. IVTT mixtures contained 2 U/μL Recombinant RNase Inhibitor ver.2 (TaKaRa). The leftmost gray bar indicates the standard IVTT condition without polymerase supplementation. T7Pol-v2 is T7 RNA Polymerase ver.2 (TaKaRa), ExpMax is Enzyme Mix in T7 RiboMAX™ Express Large Scale RNA Production System (Promega), ScriptMAX is Thermo T7 RNA Polymerase in ScriptMAX® Thermo T7 Transcription Kit (Toyobo), Hi-T7 is Hi-T7® RNA Polymerase (New England Biolabs), and HiScribe is T7 RNA Polymerase Mix in HiScribe® T7 High Yield RNA Synthesis Kit (New England Biolabs).

**Figure S4.**
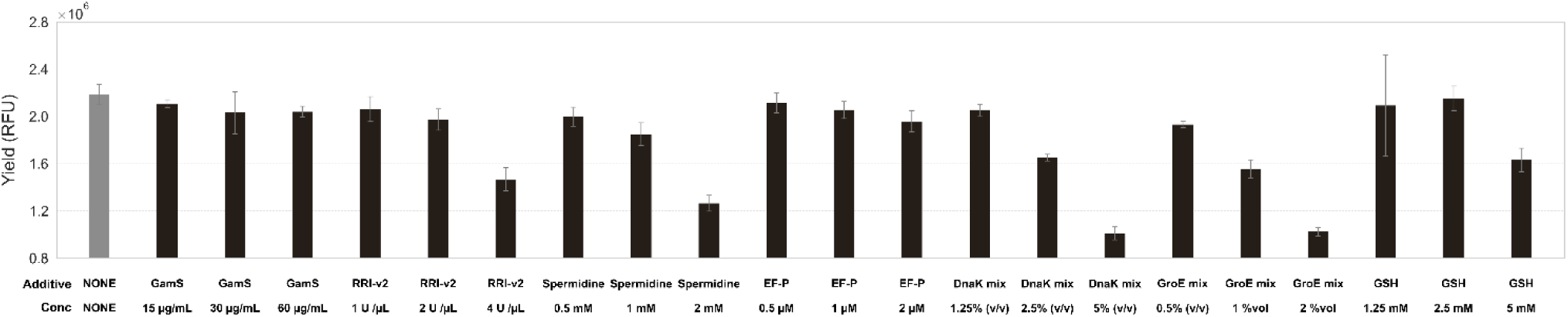
Titration of potential IVTT enhancers to the low-DNA-optimized PUREfrex1.0. mNG yields at 20 h in the low-DNA-optimized condition (T7Pol-v2 = 15% (v/v), Ribosome = 400 nM) supplemented with potential IVTT enhancers. Error bars indicate standard deviations (n = 3). The leftmost gray bar indicates the standard IVTT condition. GamS is NEBExpress® GamS Nuclease Inhibitor (New England Biolabs), RRI-v2 is Recombinant RNase Inhibitor ver.2 (TaKaRa). Spermidine was purchased from Sigma. EF-P, DnaK mix, and GroE mix were purchased from GeneFrontier. GSH was purchased from Nacalai Tesque.

**Figure S5.**
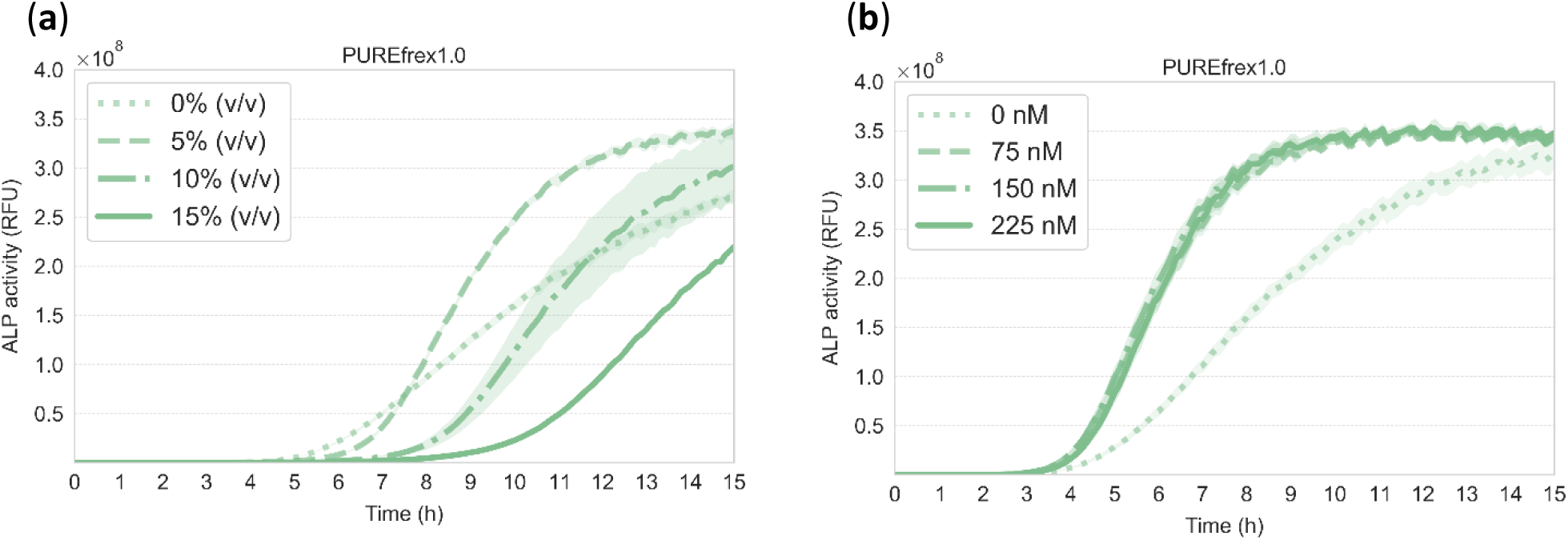
Enhancement of ALP IVTT required buffer exchange of T7 RNA polymerase ver.2. Kinetic profiles of ALP signals upon dephosphorylation of Fluorescein diphosphate at DNA = 2 pM and Ribosome = 400 nM in PUREfrex1.0. Concentrations of supplemented T7 RNA polymerase ver.2 are shown in the legends. (**a**) T7 RNA polymerase ver.2 was supplemented without buffer exchange. Signal increase in ALP activity was delayed in a dose-dependent manner. (**b**) Buffer-exchanged T7 RNA polymerase ver.2 was supplemented. Shaded areas indicate standard deviations (n = 3).

**Figure S6.**
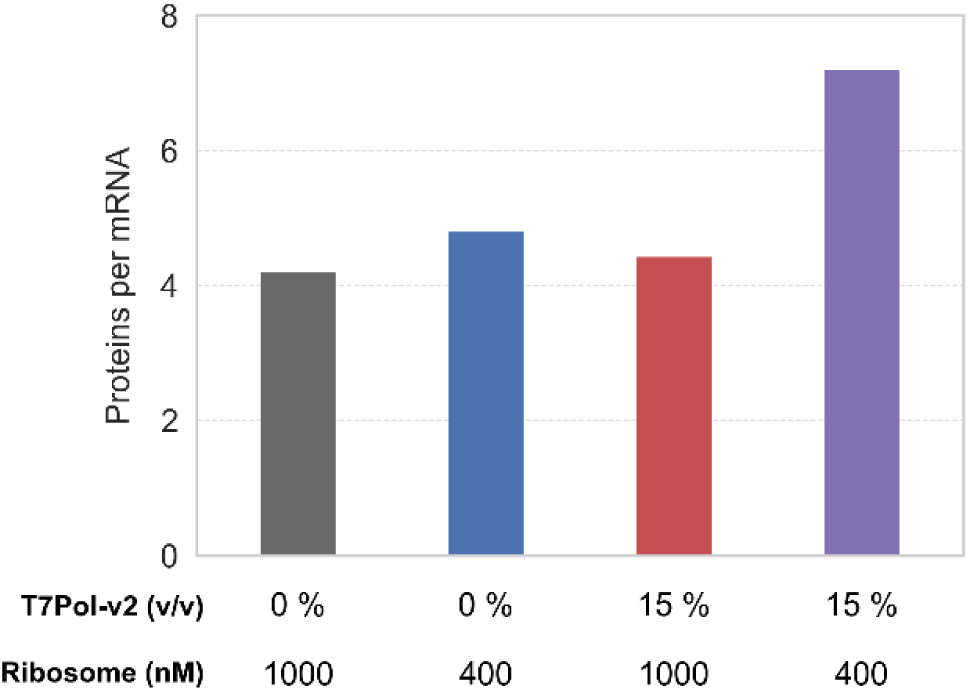
Enhanced mNG protein production per mRNA in the low-DNA-optimized IVTT. The ratios of mNG proteins over mRNA at 18 h in Figure 4c-d were plotted.

**Figure S7.**
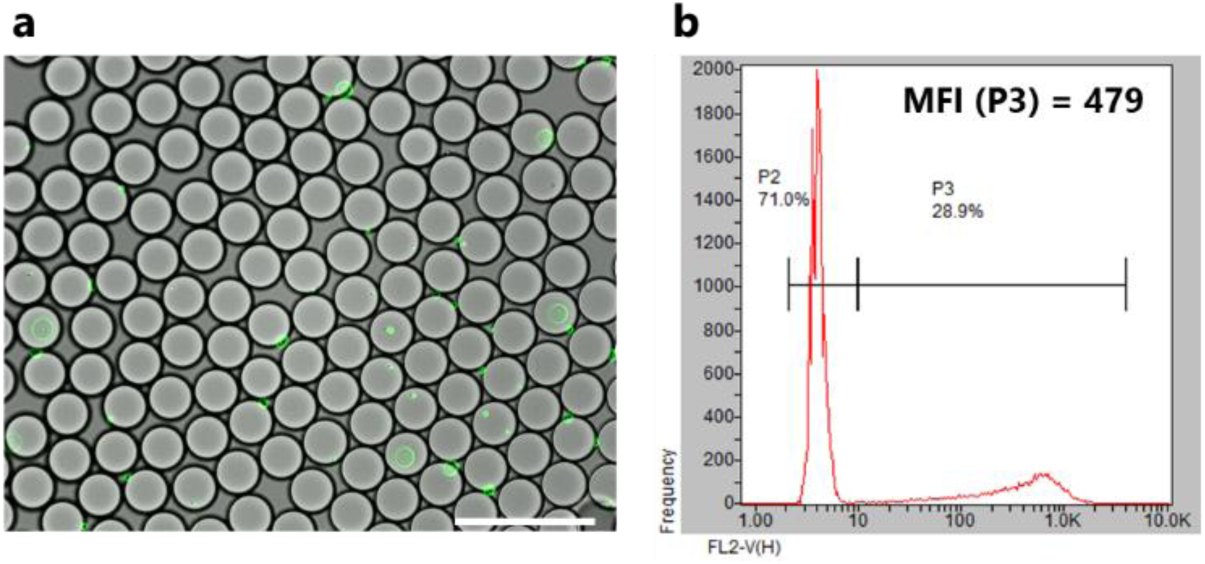
Microdroplets encapsulating *E. coli* cells expressing mNG. **(a)** Fluorescent microscopy images of droplets encapsulating *E. coli* cells expressing mNG. A scale bar is 100 μm. (**b**) Fluorescence-activated droplet analysis of droplets encapsulating *E. coli* cells expressing mNG. Mean fluorescent intensity of droplets containing *E. coli* cells was shown.

